# Modeling Spectroradiometric Measurements of Oral Mucosal Autofluorescence

**DOI:** 10.1101/2025.06.19.660587

**Authors:** Joyce E. Farrell, Xi Mou, Brian A. Wandell

## Abstract

This study explores the potential of quantitative autofluorescence imaging (AFI) as an objective tool for monitoring the health of oral mucosal tissue. Our approach involves acquiring spectroradiometric measurements of tissue fluorescence and utilizing a model to understand the fluorophores influencing these measurements, including the impact of blood attenuation. We acquired fluorescence measurements from the dorsal tongue and inner lip of healthy human volunteers, subsequently fitting the model to these data to estimate individual fluorophore contributions and the optical density of the blood. Our dataset and model, which are freely shared in an open repository, aim to advance the development of quantitative, non-invasive diagnostic imaging systems for monitoring oral health, ultimately facilitating the detection and characterization of oral mucosal tissue abnormalities.

## Introduction

Tissue autofluorescence is a phenomenon in which certain molecules within the tissue absorb light at specific wavelengths and then re-emit light at longer wavelengths. This property can be used as a non-invasive tool for monitoring tissue health. The autofluorescence signal originates from several types of fluorophores. The fluorophores NADH (nicotinamide adenine dinucleotide) and FAD (flavin adenine dinucleotide) are involved in cellular metabolism in the tissue’s epithelial layer. Collagen and elastin are fluorophores found in the tissue’s stroma layer, where they help maintain the tissue’s strength and structure. Thus, changes in the intensity, spectral characteristics, or spatial distribution of autofluorescence may reflect changes in cellular metabolic states and/or the tissue’s structural integrity.

The sensitivity of autofluorescence to biochemical and structural changes in tissues makes it a promising candidate for developing rapid, cost-effective, and non-invasive diagnostic tools across various medical fields. These changes may be indicative of pathological conditions, including inflammation and cancer that develop over time (Richards-Kortum and Sevick-Muraca 1996; De Veld et al. 2005). Non-invasive assessment of tissue autofluorescence over time has the potential to serve as an inexpensive component of clinical practice, assisting in the monitoring of tissue health and early disease detection. Autofluorescence measurements can also be valuable in delineating tumor margins during surgical procedures (Kikuta et al. 2018).

Numerous studies report that dysplastic and cancerous lesions exhibit lower tissue fluorescence in wavelengths ranging between 500 and 600 nm (“green”), compared to the surrounding healthy tissue (Gillenwater et al. 1998; Betz et al. 1999; De Veld et al. Sep-Oct 2004, 2005; Poh et al. 2006). This observation is consistent with the hypothesis that a reduction in tissue autofluorescence signals a change in cell metabolism and/or the structural integrity of the underlying tissue. This discovery formed the basis for developing autofluorescence imaging (AFI) systems to visualize these changes (Onizawa et al. 2003; Svistun et al. 2004; Lane et al. 2006).

### Current instrumentation

Under normal viewing conditions, tissue autofluorescence in healthy oral mucosa is not visible to the naked eye. The fluorescence signal generated by endogenous fluorophores is significantly lower than that of the light reflected from the tissue. Therefore, specialized optical instrumentation is necessary to isolate and effectively detect the weak fluorescence emission signal. AFI systems have two important components that make it possible to measure tissue fluorescence. First, they include an intense short wavelength light to excite the tissue fluorophores that are present in oral mucosal tissue. Second, they embed a longpass filter in a viewing apparatus, such as a pair of goggles or an eyepiece, to block reflected light from reaching the viewer, permitting only the emitted fluorescence to pass. Clinicians are instructed to search for areas within the oral cavity that appear unusually dark when using AFI viewing devices, because dysplastic and/or cancerous tissue are thought to emit less fluorescence than healthy tissue(E. C. Yang et al. 2018),

Current commercial AFI systems are engineered to provide images that clinicians can view, but they do not measure quantitative data. The subjective interpretation of fluorescence levels depends heavily on the operator’s experience and training. The absence of quantitative outputs undermines diagnostic accuracy, exacerbates overdiagnosis, and limits clinical adoption.

There remains a need for a non-invasive imaging system that can provide quantitative and reproducible measurements of tissue autofluorescence. While researchers have advanced the field by integrating autofluorescence imaging with digital cameras (Kikuta et al. 2018; Uthoff et al. 2019; Schwarz et al., n.d.; Taguchi et al. 2022)—replacing subjective clinical assessments—key challenges still persist. These include weak tissue fluorescence signals, the challenge of optimizing excitation light sources, limited spectral channels in commercial cameras, uncalibrated device components, and a lack of robust algorithms for detecting diverse oral mucosal abnormalities.

### Models and measurements

To design an AFI system that addresses these challenges, it is important to characterize the spectral signal that is available to measure. In this paper, we report on spectroradiometric measurements of light emitted and reflected from two oral regions in healthy volunteers: dorsal tongue and inner lip mucosa. We focus on the image systems components, lights and filters, that affect the generation and measurement of oral mucosal tissue autofluorescence. We specifically evaluated how the excitation light wavelength, bandwidth and intensity affect the spectral fluorescence signal.

Second, we developed a computational model that predicts the tissue fluorescence spectral signal. The model uses a linear combination of spectral basis functions representing the effects of fluorophore emissions along with a model of blood absorption. For the inner lip mucosa, fluorescence was represented by two primary components: collagen (adjusted for blood absorption) and FAD. Modeling dorsal tongue mucosa required 3–5 components, including blood-attenuated collagen, FAD, keratin, and—depending on the individual—porphyrins (present in oral bacteria) and/or chlorophyllA (linked to dietary residues).

### Related literature

There is a large literature on endogenous and exogenous fluorophores that are present in the oral cavity and measured in a variety of ways. We delay our review of the literature to the Discussion, where we quantitatively compare our measurements to multiple published reports.

## Methods

### Subjects

Four healthy adult volunteers (three males, one female; ages 30 to 73) participated in this study. Participants included two of the authors and two members of the research lab. Participants had no history of oral disease or recent antibiotic use. All participants provided written informed consent prior to participation, and the study protocol was approved by the Stanford University Institutional Review Board (IRB protocol 80769). The study was conducted in a laboratory setting, and all data were anonymized to protect participant confidentiality.

### Excitation lights

We acquired spectroradiometric measurements by illuminating two different regions of the oral cavity in 4 healthy individuals: the dorsal surface of the tongue and the mucosal tissue of the lower inner lip area. Each area was illuminated using one of three different excitation light sources.

Figure 1A illustrates the positions of the excitation lights, the spectroradiometer (SpectraScan® Spectroradiometer PR-670), and the diffusely reflecting white reference target (SENSING Standard White Target A002, 37101900197). The excitation lights were generated by combining LEDs with optics that produced a beam angle with a full width at half maximum (FWHM) of 16 degrees. There were three LEDs, with nominal peak wavelengths at 405 nm, 415 nm, and 450 nm. A 450 nm short-pass filter was positioned in front of the excitation lights and the illuminated surfaces, all set at a distance of 10 cm. This filter effectively reduced the spectral energy at wavelengths above 450 nm. Measurements were performed in a dark room with black-painted walls.

**Figure 1.**
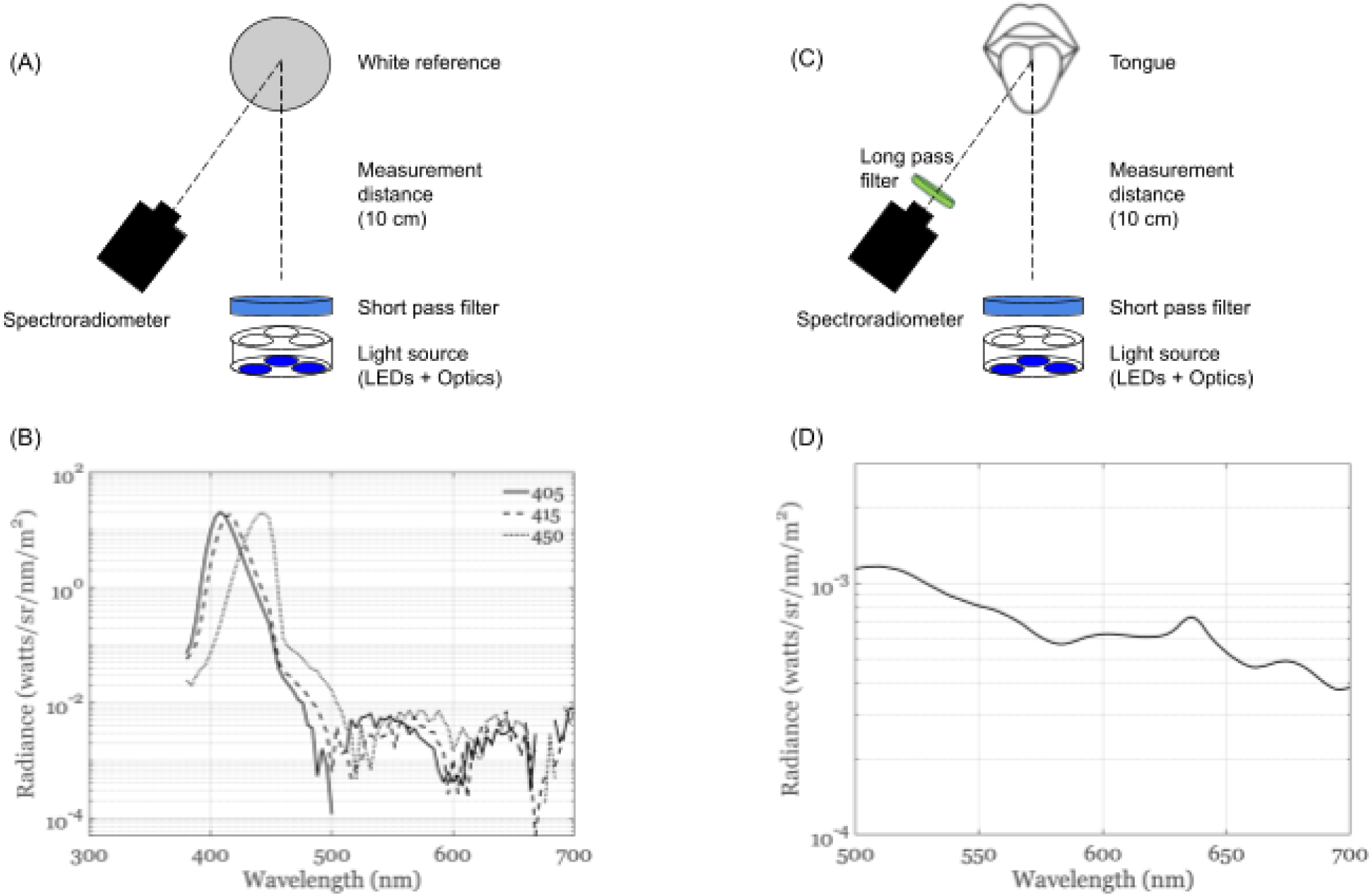
**A, B: Geometry and filters for the light measurement.** These panels depict the measurement of the excitation lights. The nominal wavelengths for the lights are 405 nm, 415 nm, and 450 nm, but their actual peak wavelengths are 408 nm, 416 nm, and 444 nm, respectively. To minimize unwanted long-wavelength light, the excitation lights pass through a 450 nm short-pass filter. The graph in panel B displays the spectral radiance of these excitation lights on a logarithmic scale. **C, D: Geometry and filters for the oral cavity measurement.** These panels show the setup for measuring light from the oral cavity. This arrangement is similar to the light measurement setup, but a 475 nm long-pass filter is positioned in front of the spectroradiometer. The graph in panel D shows the spectral fluorescence emitted from the dorsal tongue when illuminated with a 415 nm excitation light, also on a logarithmic scale. Note that the measured wavelength range for fluorescence is 500-700 nm, and the fluorescence signal is approximately four orders of magnitude lower than the peak radiance of the excitation light. The 475 nm long-pass filter is crucial for blocking reflected excitation light, which enables the long exposure time needed to detect the weak fluorescent signal.

Figure 1B displays the radiance of the three excitation lights. The intensity of each light was precisely controlled by adjusting the input current to the LEDs, which operated in constant current mode with regulation to within less than 0.01%. Each light was allowed to reach thermal equilibrium and stable luminous output before measurements were taken. The lights were calibrated by measuring the light reflected from a diffusive reflectance white reference target, using a PR-670 spectroradiometer positioned approximately 1 meter from the illuminated surface at an angle of 45 degree from the incident light axis. The reference white target surface was a diffusely reflective (Lambertian) material, for which luminance remains constant regardless of the observation distance or angle, assuming a uniform illumination. Although the measurement distance between the spectroradiometer and the reference target is not critical due to the Lambertian nature of the reflective surface, all measurements were conducted under consistent geometric and instrumental settings to ensure repeatability. The PR-670 instrument measures at 4 nm sampling intervals.

Figure 1C shows the configuration for measuring the oral cavity. In this case, a 475 nm long pass filter was placed in front of the spectroradiometer. This was necessary to enable the instrument to capture the relatively weak fluorescence signal, blocking the very bright reflected light from reaching the spectrophotometer. Figure 1D is an example measurement.

The distance from the oral cavity to the excitation light was about 10 cm and the distance from the oral cavity to the PR670 was about 1 m. Hence, the spectroradiometer measured the fluorescence from a circular area about 1.7 cm in diameter.

### Tissue fluorescence levels

For each spectral measurement, subjects were instructed to keep their tongue or lower lip steady while the intensity of the excitation light was adjusted and spectral radiance was recorded. Subjects wore safety goggles that blocked the excitation light and rest periods were provided between measurements at different excitation wavelengths (405, 415, and 450 nm) and intensities to minimize muscle fatigue.

Reflected light was estimated from two measurements. First, we measured the amount of long wavelength excitation light that escaped through the 475 nm low pass filter. (Beyond 680 nm, the reflected light from the white surface was either absent or too faint for the PR-670 to measure accurately.) Second, we measured tissue reflectance by illuminating the dorsal tongue and inner lip regions with a broadband tungsten light and dividing the measured radiance with the radiance of the reference white target that was also illuminated by the same light. Multiplying the tissue reflectance with the excitation light estimates the amount of light reflected from tissue that could potentially interfere with measurements of tissue fluorescence. Across all conditions and subjects, the tissue fluorescence level exceeded the unwanted reflected light level by about an order of magnitude (Figure 2).

**Figure 2.**
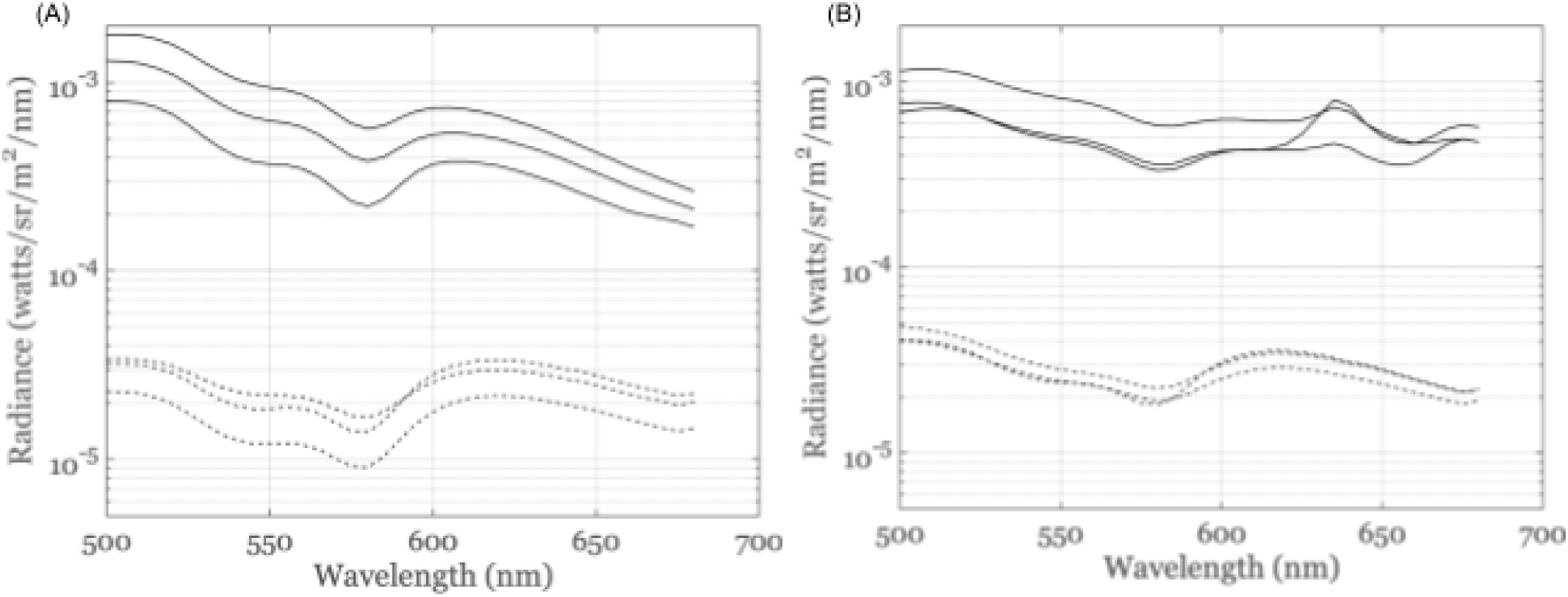
Fluorescence signal compared to unwanted reflected light for a 415 nm excitation light. The solid curves are fluorescent signals. The dashed curves are estimates of the reflected light. The symbols indicate the subject. (A) Lip. (B) Tongue. The data are plotted on a logarithmic (log₁₀) scale to highlight the fact that the tissue fluorescence levels exceeds the unwanted reflected light by an order of magnitude. Both tissue fluorescence and reflectance show evidence of blood absorption with dips at 540 and 580 nm.

## Results

### Data and software sharing

We provide freely available and open source GitHub repositories for this paper^1^, ISETCam^2^, and isetfluorescence^3^ that can be used to reproduce all the analyses and figures in this paper. The repository contains (a) the spectroradiometric measurement data, (b) the analysis software, and (c) the scripts that generate the figures in this paper.

### Spectral fluorescence from the dorsal tongue and inner lip

The amplitude of the fluorescence signals measured from the dorsal tongue and the inner lip increased with the intensity of the excitation light. After normalizing the spectra to 1 at 520 nm, the curves for each excitation light level and tissue type closely overlap in the 500 to 600 nm range (Figure 3). This indicates that for all subjects, within this spectral band, the shape of the fluorescence spectra remains consistent within each tissue type, regardless of the excitation light intensity.

**Figure 3.**
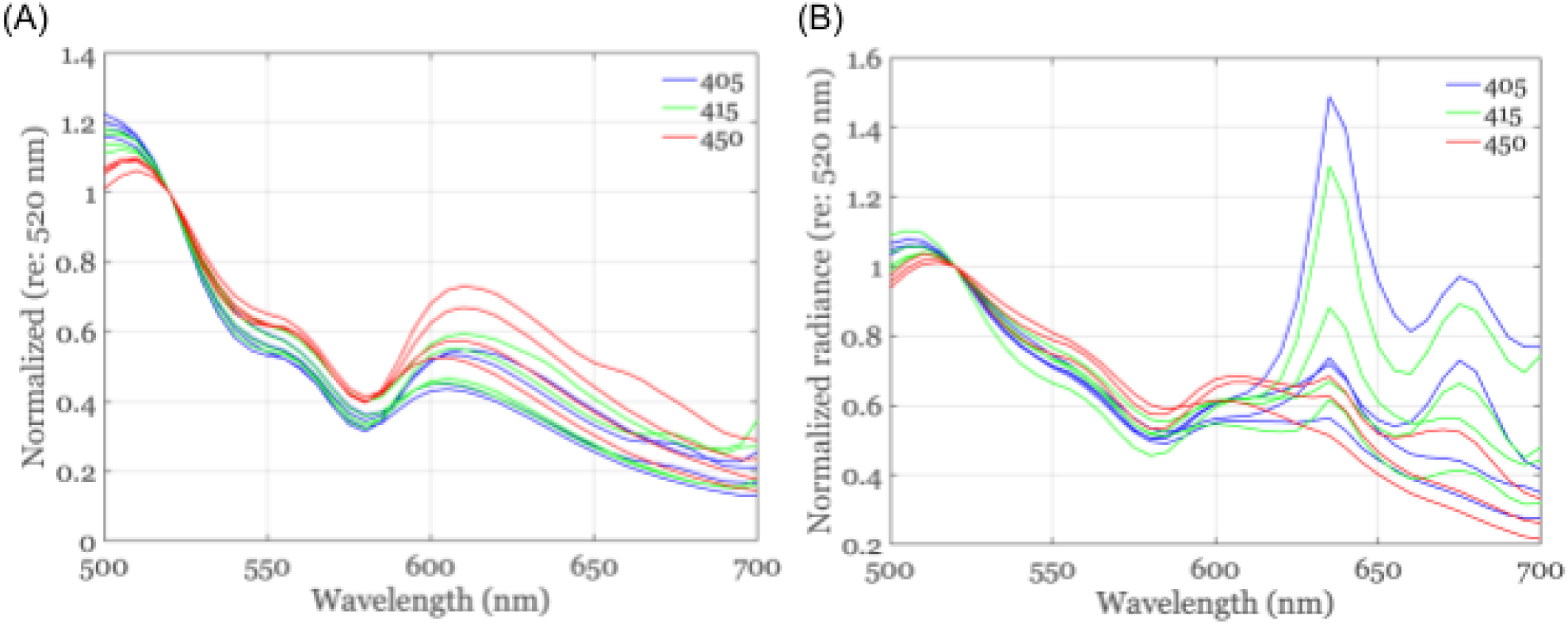
Fluorescence measured with the three different excitation lights. Lines of the same color used the same excitation light; within a color each curve is from a different subject. (A) Lip fluorescence. The data are very similar, although there is more overlap in the fluorescence using 405 nm and 415 nm excitation lights. (B) Tongue fluorescence. The data are very similar in the 500-600 nm, again with more overlap for the 405 nm and 415 nm excitation light. However, there is more variation between subjects in the fluorescence measured in the 600 - 700 nm band, which we explain in the analysis. All curves are scaled to 1 at 520 nm.

### Classifying by excitation light

For the inner lip, illumination with 405 nm and 415 nm excitation lights produced very similar fluorescence spectra for all subjects throughout the 500 nm to 700 nm measurement range. Measurements using the 450 nm excitation light, however, are systematically higher (Figure 3a). Therefore, for subsequent analysis and modeling, data from the 405 nm and 415 nm excitation lights are combined, while data from the 450 nm excitation light are analyzed separately. For the dorsal tongue, there are large signals and individual differences in the fluorescence spectra between 600 nm and 700 nm, but not when exciting with the 450 nm light. This supports our decision to analyze the 405/415nm data together and the 450 nm data separately (Figure 3b).

### Classifying by tissue type

The differences between the spectral signals of the two tissue types are further illustrated in Figure 4, which displays data from all subjects, collected across multiple days, excitation intensities, and various locations on the dorsal tongue (solid lines) and inner lip (dashed lines). Panel A presents measurements with the 405/415 nm excitation lights, while Panel B shows results with the 450 nm excitation light. The measurements from the lip and tongue separate into different clusters, for both excitation lights.

**Figure 4.**
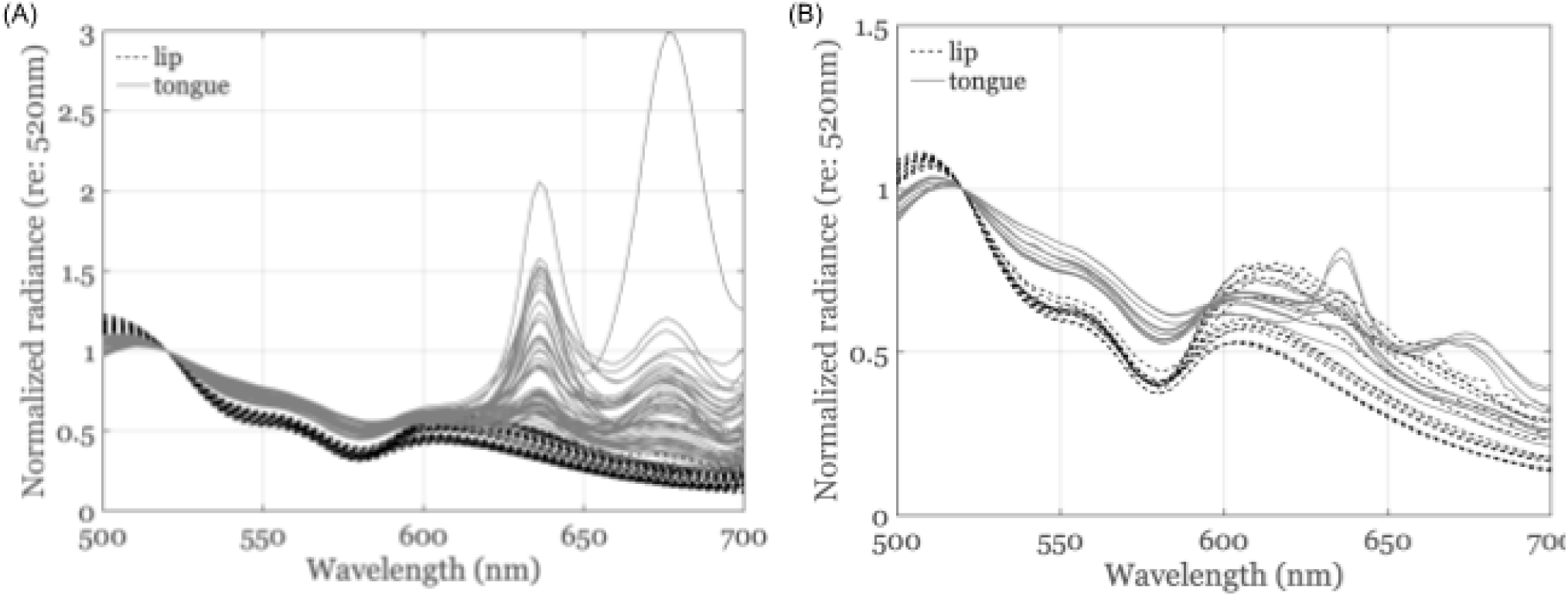
Spectral radiance of dorsal tongue and inner lip oral mucosal tissue measured with excitation lights at (A) 405/415 nm and (B) 450 nm. The data were acquired from four subjects on different days, different excitation light intensities, and different locations on the dorsal tongue (solid line) or on the inner lip (dashed line). The spectral radiance curves are scaled to 1 at 520 nm.

### Modeling

We fit the spectral fluorescence curves using a model based on (a) fluorophore emission spectra, and (b) spectral transmission through the blood. The same framework is used for fitting both tongue and lip, differing only in that the lip has a subset of the fluorophores present on the tongue.

Figure 5A shows the main tissue layers and the fluorophores that are likely to be found in each layer of the tongue and lip. Collagen is concentrated in the underlying stromal connective tissue, which also houses blood vessels and nerves. FAD and NADH are present in the cytoplasm of epithelial cells that reside in the superficial layer of oral mucosal tissue. On the tongue, these cells are sometimes covered by an additional layer of fluorescent keratinized cells. Also, some individuals have porphyrins (from bacteria or food) and chlorophyll-a (from green vegetables) on the dorsal tongue. Thus, fluorescence measured from tissue on the dorsal tongue and inner lip will depend on FAD, collagen, and blood. The dorsal tongue signal can also depend on keratin, porphyrins, and chlorophyll-a.

**Figure 5.**
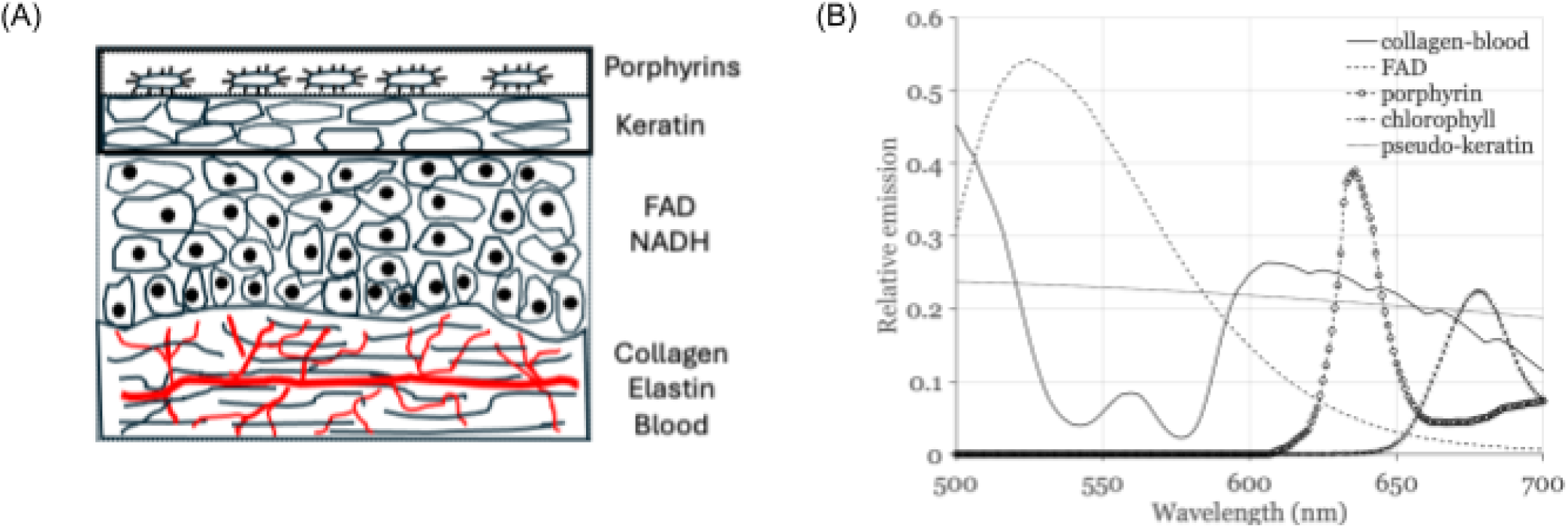
Model description. (A) Description of the spatial distribution of the fluorophores in the tongue and lip. FAD, NADH, Collagen and blood are present in both. The tongue has an additional layer of cells with keratin, and the surface may contain both porphyrin and chlorophyll-a. The chlorophyll-a can be present from residual food, such as leafy vegetables. (B) The spectral emissions of the fluorophores. The collagen is illustrated after transmission through the blood. Filtering by the blood creates the local minima near 540 and 577 nm.

The spectral emission curves of the fluorophores are shown in Figure 5B. The emission spectra of collagen and FAD are based on excitation-emission matrices from DaCosta et al 2003. We plot the collagen emissions as they would be measured after passing through blood^4^ (Figure 5A), which we model as a spectral filter. Porphyrins and chlorophyll-a have unique and identifiable emission spectra and their concentration depends on the diet of the participant.

There is consensus that keratin emission spectra are broad and overlap significantly with FAD, but there is little agreement about the precise keratin emission spectra (Malak et al. 2022). Blood concentration can also vary between participants and oral cavity locations. For this reason we estimate the keratin emission spectrum and blood concentration as part of the fitting process (see below).

We do not model NADH or elastin. We eliminate NADH because its excitation sensitivity does not extend to 405/415 and 450nm (Georgakoudi et al. 2002). We exclude elastin because it has relatively low concentration in the stromal layer compared to collagen (Pavlova et al. 2008; Banerjee, Miedema, and Chandrasekhar 1999).

### Numerical fitting procedure

We model the total fluorescence measured from a tissue area (**bulk fluorescence**) as the weighted sum of the emissions from various fluorophores. The model accounts for the fact that collagen, but not the other fluorophores, is transmitted through blood. The impact of blood is important, and many studies report dips in oral mucosal tissue fluorescence around 540 and 577 nm, where there is significant oxygenated blood absorption (De Veld et al. 2005). Because oxyhemoglobin is generally present in higher concentration than deoxyhemoglobin, we model only oxygenated blood.

We fit the data in two parts. First, we perform a preprocessing step to estimate keratin’s spectral emission. We then use the estimated keratin emission to fit the data from each individual.

### Estimation procedures

We model the measured spectral bulk fluorescence, F(λ), as the weighted sum of N fluorophores:

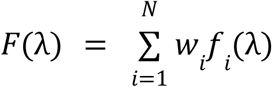

There is one special consideration: the first fluorophore, *f*_1_ (λ), is always the emission spectrum of collagen filtered through the blood at a certain optical density (*d*) that we estimate. The optical density represents the degree to which light is absorbed by the blood.

The spectral transmission through the blood is calculated from the blood optical density. Given an optical density, *d*, and an oxyhemoglobin absorbance *a*(λ), the spectral transmittance is calculated as:

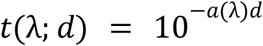

The spectral absorbance of blood is based on data provided by (Prahl, n.d.). The collagen emission spectrum, *c*(λ), is provided in (DaCosta, Andersson, and Wilson 2003) and plotted in Figure 5 in this paper. Therefore, the first term in the sum, *f*_1_ (λ), representing the collagen emission filtered through blood, is calculated as:

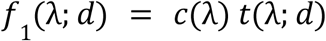

We can rewrite the model more explicitly as:

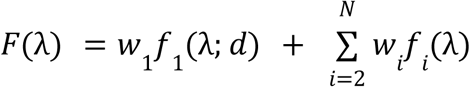

We solve for the optical density, *d*, and the weights, *w_i_*, using a non-negative least squares method.

### Estimation of keratin’s spectral emission

We needed to perform one preprocessing step to estimate the unknown keratin emission spectrum. We modeled keratin as a skewed Gaussian function of wavelength:

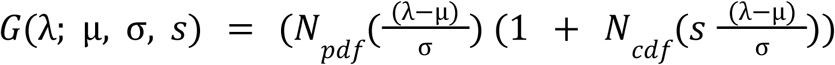

where:

- *N*_*pdf*_ is the Normal probability density function.
- *N*_*cdf*_ is the Normal cumulative density function.
- µ, σ, *s* are the mean, standard deviation, and skewness parameters of the skewed Gaussian function, respectively.

Using the tongue spectral fluorescence data from all the subjects, we performed a single global fit for blood density, the three unknown keratin parameters, and we allowed the fluorophore weights to vary between measurements. Only the keratin parameter values derived from this search were reused in the subsequent fits to the data from individual subjects. The fifth fluorophore, pseudo-keratin, is estimated as:

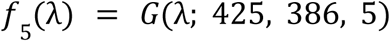

Figure 5B illustrates the resulting pseudo-keratin emission and the blood-attenuated collagen spectrum.

After the keratin model was established, we estimated the blood density and fluorophore parameter weights again for each bulk fluorescence measurement. For the tongue, we fit each curve with a search that finds the non-negative weights of the five fluorophores (collagen, FAD, porphyrin, chlorophyll-A, keratin) and one parameter that estimates the blood density for each individual. For the lip, the final estimates use only two fluorophores (collagen and FAD), and one parameter that estimates the blood density for each individual^5^.

The blood density and fluorophore parameters are expected to depend on the wavelength of the excitation light and the local tissue properties. The location of the blood is below the surface of the tissue and the excitation light will penetrate a different amount, depending on wavelength. The fluorophores have wavelength-dependent excitation functions, so their weights too should depend on the excitation light wavelength.

### Bulk fluorescence fits: 405/415 nm excitations

#### Tongue spectral fluorescence

Each of the four panels in Figure 6 displays the tongue bulk fluorescence measurements for an individual subject, along with the corresponding model fit (excitation 405/415 nm). The large impact of porphyrin (peak 636 nm) and chlorophyll-a (peak 675 nm) varies greatly between participants and is obvious in the data. The similarity between all participants in the 500 - 600 nm band, along with the local minima arising from the blood transmission, is also captured. We provide the estimated weights of the different fluorophores and blood optical density in Table 1.

**Figure 6.**
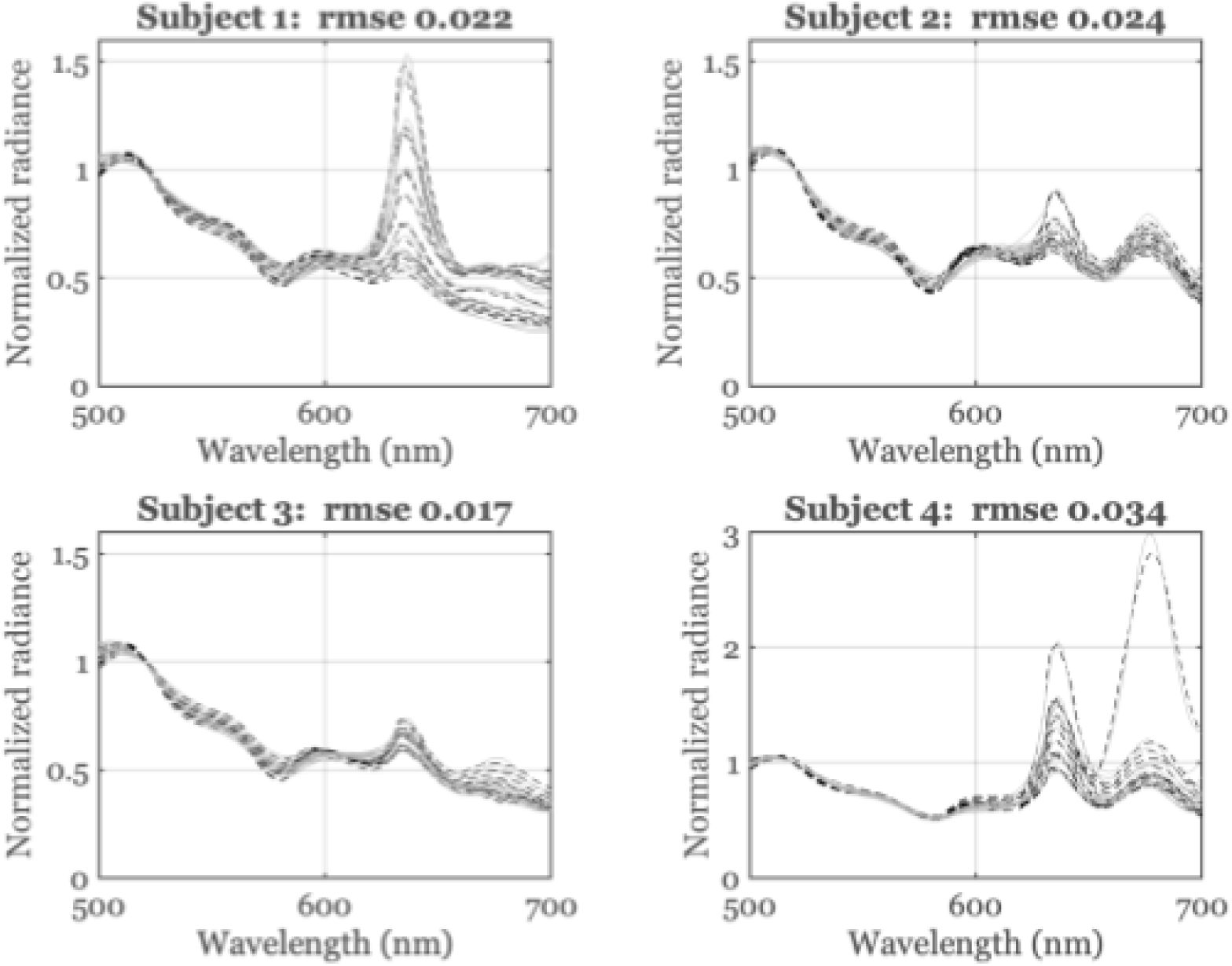
Tongue bulk fluorescence from the 405/415 nm excitation lights. Each panel shows a different subject. The data include different excitation light intensities, different days and locations. The gray shaded curves are measurements, scaled to 1 at 520 nm, and the dashed black curves are model fits.

**Table 1.** Subject-specific estimates of blood optical density and fluorophore weight for the tongue. Each row lists the estimated values for blood optical density and mean fluorophore weight for each subject. The number of measurements (N) that were used for each estimate are indicated in parentheses. Details on the calculation methods are provided in the main text

| Parameter | Excitation light | Subject 1 | Subject 2 | Subject 3 | Subject 4 |
| --- | --- | --- | --- | --- | --- |
| **Blood density** | 405/415 | 1.00<br>(N = 15) | 1.07<br>(N = 14) | 0.98<br>(N = 14) | 1.12<br>(N = 15) |
|  | 450 | 1.24<br>(N = 3) | 1.25<br>(N = 3) | 1.23<br>(N = 3) | 1.22<br>(N = 2) |
| **Collagen** | 405/415 | 8.69 ± 1.07<br>(N = 15) | 13.18 ± 1.05<br>(N = 14) | 9.23 / 1.16<br>(N = 14) | 10.72 / 0.77<br>(N = 15) |
|  | 450 | 8.90 ± 0.44<br>(N = 3) | 12.77 ± 1.87 | 13.83 ± 0.45<br>(N = 3) | 10.21 / 1.38<br>(N = 2) |
| **FAD** | 405/415 | 0.55 / 0.05<br>(N = 15) | 0.55 ± 0.02<br>(N = 14) | 0.54 ± 0.02<br>(N = 14) | 0.52 ± 0.05<br>(N = 15) |
|  | 450 | 0.72 ± 0.02<br>(N = 3) | 0.60 ± 0.00<br>(N = 3) | 0.71 ± 0.01<br>(N = 3) | 0.61 ± 0.03<br>(N = 2) |
| **Porphyrins** | 405/415 | 0.42 ± 0.29<br>(N = 15) | 0.20 ± 0.09<br>(N = 14) | 0.19 ± 0.04<br>(N = 14) | 0.73 ± 0.30<br>(N = 15) |
|  | 450 | 0.05 ± 0.02<br>(N = 3) | 0.06 ± 0.05<br>(N = 3) | 0.09 ± 0.01<br>(N = 3) | 0.24 ± 0.05<br>(N = 2) |
| **Chlorophyll-a** | 405/415 | 0.04 ± 0.04<br>(N = 15) | 0.27 ± 0.04<br>(N = 14) | 0.04 ± 0.04<br>(N = 14) | 0.54 ± 0.51<br>(N = 15) |
|  | 450 | 0.00 ± 0.00<br>(N = 3) | 0.06 ± 0.05<br>(N = 3) | 0.00 ± 0.00<br>(N = 3) | 0.09 ± 0.02<br>(N = 2) |
| **Keratin** | 405/415 | 0.32 ± 0.08<br>(N = 15) | 0.23 ± 0.03<br>(N = 14) | 0.30 ± 0.05<br>(N = 14) | 0.36 ± 0.04<br>(N = 15) |
|  | 450 | 0.30 ± 0.04<br>(N = 3) | 0.35 ± 0.03<br>(N = 3) | 0.20 ± 0.00<br>(N = 3) | 0.35 ± 0.01<br>(N = 2) |

#### Inner lip spectral fluorescence

Figure 7 shows the measurements and model fit to the lip bulk fluorescence for each of the four subjects (excitation 405/415 nm), plotted as four separate panels. The fits exclude porphyrin and chlorophyll-a, and the spectral emissions agree across the entire 500 - 700 nm range. The similarity between all participants, along with the local minima arising from the blood transmission, is also captured. We include a table of the estimated fluorophore and blood optical density weights in Table 2.

**Figure 7.**
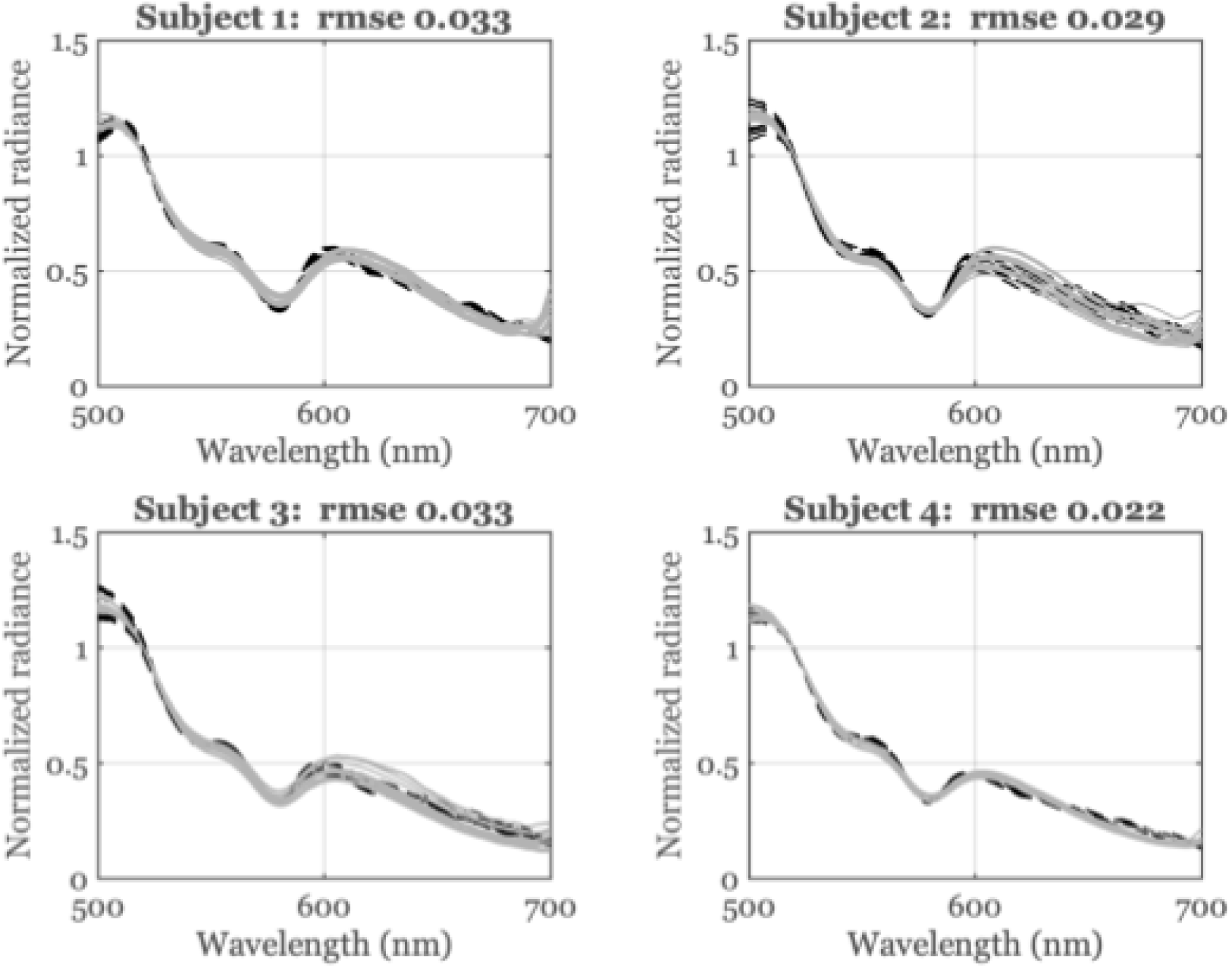
Lip bulk fluorescence from the 405/415 nm excitation lights. Each panel shows a different subject. The curves include different excitation light intensities, different days and locations. The gray shaded curves are measurements, scaled to 1 at 520 nm, and the dashed black curves are model fits.

**Table 2.** Subject-specific estimates of blood optical density and fluorophore weight for the lip. Each row lists the estimated values for blood optical density and mean fluorophore weight for each subject. The number of measurements (N) that were used for each estimate are indicated in parentheses. Details on the calculation methods are provided in the main text

| Parameter | Excitation light | Subject 1 | Subject 2 | Subject 3 | Subject 4 |
| --- | --- | --- | --- | --- | --- |
| **Blood density** | 405/415 | 1.06<br>(N=14) | 0.96<br>(N = 13) | 0.80<br>(N=14) | 0.78<br>(N=14) |
|  | 450 | 1.24<br>(N=3) | 1.25<br>(N=3) | 1.02<br>(N=3) | 1.10<br>(N=4) |
| **Collagen** | 405/415 | 17.34 ± 0.52<br>(N=14) | 16.12 ± 1.41<br>(N = 13) | 13.12 ± 0.93<br>(N=14) | 11.81 ± 0.38<br>(N=14) |
|  | 450 | 22.18 ± 1.81<br>(N=3) | 24.61 ± 0.97<br>(N=3) | 14.58 ± 0.99<br>(N=3) | 15.19 ± 1.21<br>(N=4) |
| **FAD** | 405/415 | 0.63 ± 0.01<br>(N=14) | 0.54 ± 0.03<br>(N = 13) | 0.49 ± 0.02<br>N=14 | 0.53 ± 0.02<br>N=14 |
|  | 450 | 0.75 ± 0.02<br>(N=3) | 0.73 ± 0.00<br>(N=3) | 0.66 ± 0.01<br>(N=3) | 0.72 ± 0.02<br>(N=4) |

### Bulk fluorescence fits: 450 nm excitation

The bulk fluorescence data and corresponding model fits for the tongue (Figure 8) and lip (Figure 9) are shown for a 450 nm excitation light. The 450 nm excitation is relatively ineffective at producing porphyrin and chlorophyll-a emissions on the tongue. Thus, the weights for these exogenous fluorophores are low for the 450 nm excitation light and the data between subjects is consistent over the whole range, from 500 - 700 nm.

**Figure 8.**
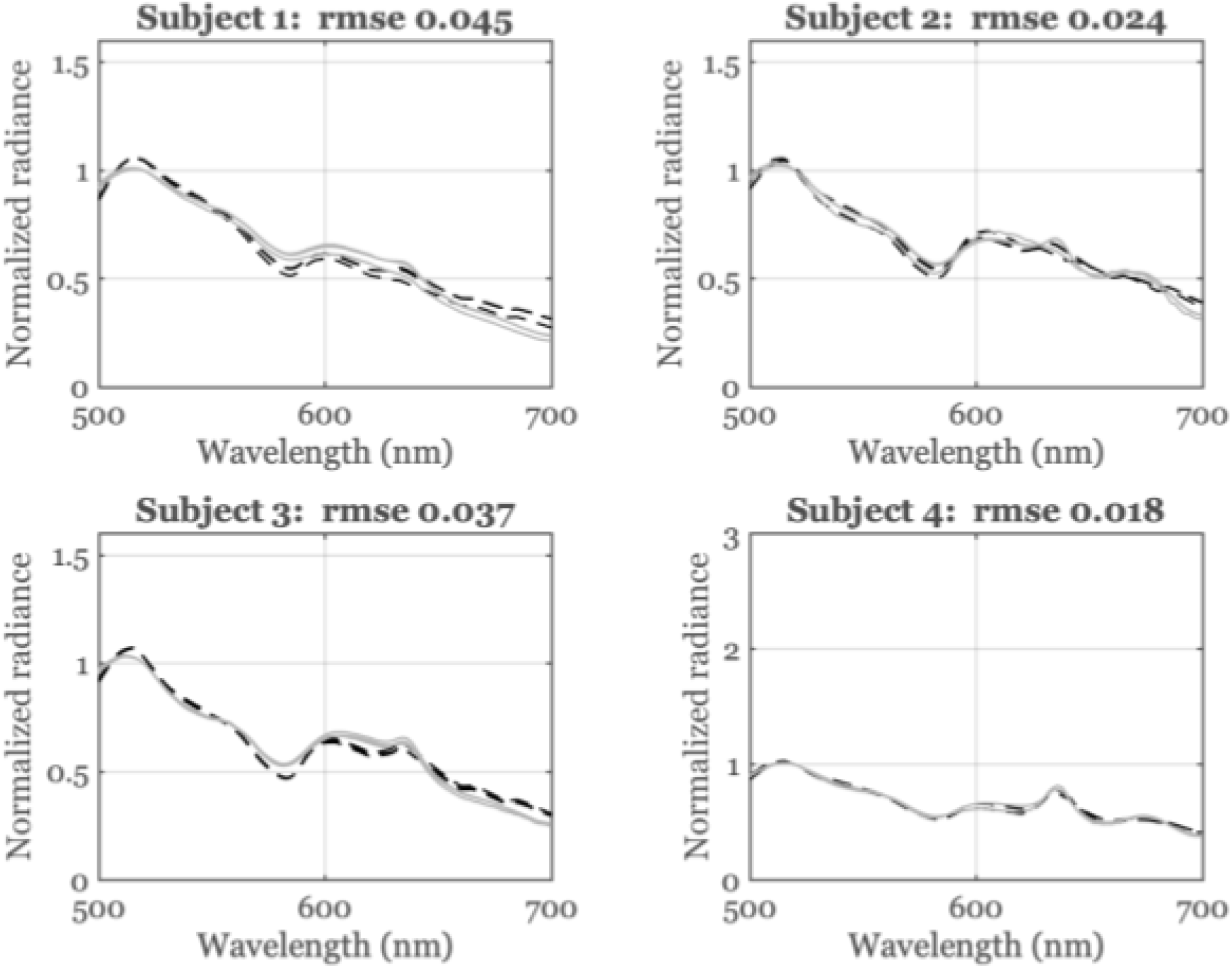
Tongue bulk fluorescence from a 450 nm excitation light. Each panel shows a different subject. The curves include different excitation light intensities, different days and locations. The gray shaded curves are measurements, scaled to 1 at 520 nm, and the dashed black curves are model fits.

**Figure 9.**
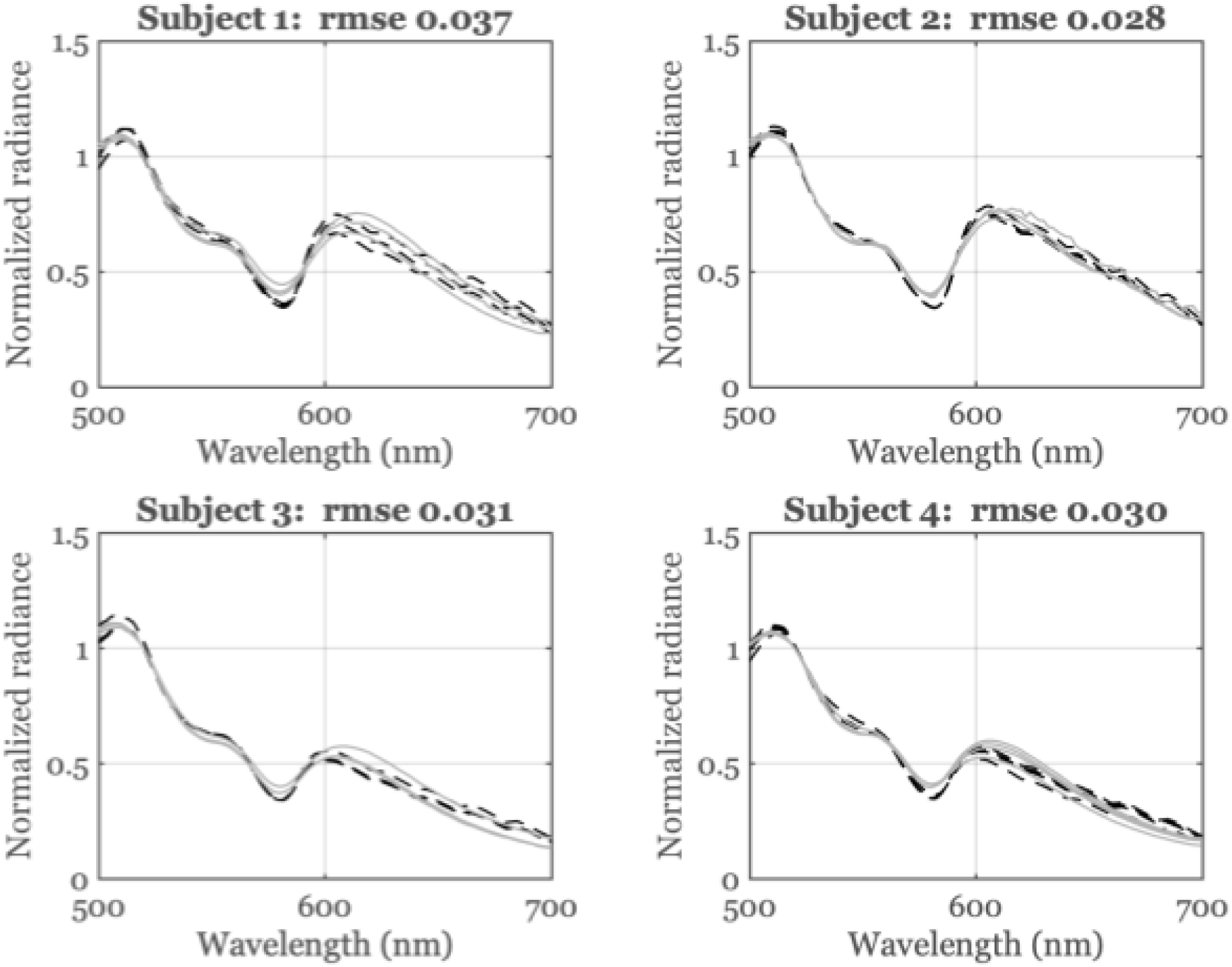
Lip bulk fluorescence and model fits from a 450 nm excitation light. Each panel shows a different subject. The curves include different excitation light intensities, different days and locations. The gray shaded curves are measurements, scaled to 1 at 520 nm, and the dashed black curves are model fits.

The local minima at the blood absorption wavelengths are more pronounced for the 450 nm excitation light than the 405/415nm light, consequently, the collagen emissions must pass through more blood. We discuss this more fully in the next section, which includes the blood optical density and fluorophore weight estimates (Tables 1 and 2).

### Model weights

Tables 1 (tongue) and 2 (lip) present subject-specific blood density and fluorophore weights. Each column in the tables corresponds to a different subject, while the rows represent the estimated values for each parameter. A single blood optical density weight was determined for each combination of (a) subject, (b) substrate (lip or tongue), and (c) excitation light (405/415 nm or 450 nm). Since multiple measurements were taken for the same subject, substrate, and excitation light, the fluorophore weights were calculated for each measurement, and we report the mean and standard deviation.

Across both tongue and lip measurements, the estimated FAD weights are consistent between subjects. Notably, for all four subjects, the FAD weight was higher when excited with 450 nm light compared to 405/415 nm excitation. Similarly, blood density values were consistently higher under 450 nm excitation than under 405/415 nm excitation for all subjects and both substrates.

Regarding collagen, the estimated weights in the lip were higher with 450 nm excitation compared to 405/415 nm. However, collagen weights in the tongue did not show a significant difference between the two excitation wavelengths.

For the tongue measurements, the estimated weights of porphyrin and chlorophyll-a exhibited substantial variability under the 405/415 nm excitation, but were uniformly low under 450 nm excitation. The estimated keratin weights were similar across subjects and for both excitation lights. We discuss these subject differences in the following section.

## Discussion

### Bulk fluorescence model

We introduce a simple model to account for the bulk fluorescence spectral measurements from the inner lip and dorsal tongue of the human oral cavity. Despite its simplicity, the model fits the bulk spectral fluorescence (500 - 700 nm) with a root mean squared error of 1-3%.

There have been other approaches to modeling the bulk fluorescence signal in the oral cavity. For example, investigators have used Diffusion (Müller et al. 2003), Photon Migration (Müller et al. 2001) and Monte Carlo models (Friebel et al. 2006; Pavlova et al. 2008)) to simulate how light propagates through oral mucosal tissue. These models are stochastic, and they can become quite complex. For example, they often incorporate an extensive list of tissue properties, such as the thickness of the tissue layers and scattering properties of the substrate.

The model we introduce is a compromise between the complex computational models and a purely data-driven model based on linear decomposition methods such as Principal Components Analysis (De Veld et al. 2003, Sep-Oct 2004) and Non-negative Matrix Factorization (S. Yang et al. 2023). Our model incorporates the blood optical density, and its formulation is based on the known biology of the oral cavity. The model is simple enough to enable estimation of the contributions of multiple fluorophores and the impact of the blood. In the one case where the fluorophore emission was uncertain, (keratin), we could estimate its spectral emissions using a method that is similar to multivariate curve resolution (MCR), which extracts component spectra and their relative concentrations directly from mixed spectral data (Müller et al. 2003).

To our knowledge, we are the first to model the absorption of collagen fluorescence by blood. Others have observed the dips and even attributed them to oxygenated hemoglobin, but they did not model these dips.

### Fluorophore weights and blood optical density

We obtain parameter values of the blood density and fluorophore weights across a range of conditions by fitting the model to the bulk fluorescence data measured from the dorsal tongue and inner lip of each individual and for each excitation light (Tables 1 and 2). These parameter values inform us about the biological substrate in each individual. The literature establishes several expectations about these parameters that we can compare with the model estimates.

Porphyrins and chlorophyll-a, which are exogenous fluorophores, were detected on the dorsal tongue in some subjects but not in others. This variation is expected, as exogenous fluorophores are not present in all individuals. Excitation at 450 nm did not produce a response from either porphyrin or chlorophyll-a. This result is also expected since the peak excitation wavelength for these fluorophores is typically around 400–420 nm (Table 1). Since exogenous fluorophores were not detected on the lip, they were not used to fit the data (Table 2).

The peak excitation wavelength for FAD is 450 nm. Hence, as expected, for both the inner lip and dorsal tongue in all four subjects, the weights associated with FAD are higher when measured with the 450 nm excitation light than the 405/415 excitation lights (Table 1).

The measured keratin weights remained relatively consistent regardless of the specific light wavelengths used for excitation or differences between the four study participants. While no previous studies have directly measured keratin’s light absorption properties in living tissue, existing research emphasizes that the connection between excitation wavelengths and keratin’s emitted light signals is complex (Zheng et al. 2008). Earlier experiments with purified keratin powder found only slight emission variations when tested under 405 nm, 425 nm, and 450 nm excitation lights (Wu and Qu 2006; S. Yang et al. 2023). Similarly, this study observed minimal differences in keratin measurements when comparing excitation wavelengths within this same 405–450 nm range.

Collagen weights were consistently higher in the inner lip than in the tongue. This difference may be due to structural differences: The stroma of the inner lip is relatively thick and is composed of a superficial layer known as the lamina propria, along with a deeper layer that contains a dense concentration of collagen fibers. In contrast, the stroma of the tongue is made up only of a thinner lamina propria, which attaches directly to the underlying muscle. Therefore, the thicker stromal layer in the inner lip contains more collagen fibers than that of the tongue (Ten Cate 1994), which may explain the higher collagen weights.

The peak excitation wavelength for collagen is close to 400 nm, hence one might expect the estimated weights for collagen to be higher for the 405/415 lights than for the 450 nm lights. However, the estimated weights for collagen in the inner lip are higher for the 450 nm light than the 405/415 nm light. One possible explanation for this finding is that photons must travel through the epithelial tissue layers before reaching the stromal layer where collagen and blood reside. Photons from a 450 nm light penetrate tissue more deeply than photons from a 405/415 nm light. The amount of collagen fluorescence is a balance between the absorption efficiency of collagen at the excitation light (higher at 405 than 450 nm) and the depth of tissue penetration by the excitation light (greater for 450 than 405 nm).

Finally, the estimated blood densities are higher for both the inner lip and dorsal tongue when excited with the 450 nm light compared to the 405/415 nm lights. This too may be due to the fact that 450 nm light penetrates tissue more deeply into the vascularized layers. In that case, we would expect the collagen emissions to pass through more oxygenated blood and thus have a higher optical density.

### Related literature

Numerous studies have reported in-vivo spectroradiometric measurements of oral mucosal tissue fluorescence in healthy individuals. For instance, De Veld and her colleagues (De Veld et al. 2003) used a fiber optic system to illuminate six different regions of the oral cavity with various excitation wavelengths, including 405 nm. We scanned the data from Figure 4 in their paper, which shows the mean bulk fluorescence spectra of healthy oral mucosa from the inner cheek and the dorsal surface of the tongue.

To check the consistency between their data and ours, we applied the following approach: if the spectral fluorescence in both datasets can be explained by a weighted sum of the same fluorophores, then we should be able to reconstruct the (De Veld et al. 2003) data using a weighted sum of our spectral measurements. The fit is excellent (Fig 10), indicating that our data are consistent with the central tendency of their extensive dataset, which comprises 9295 samples from 96 volunteers^6^.

**Figure 10.**
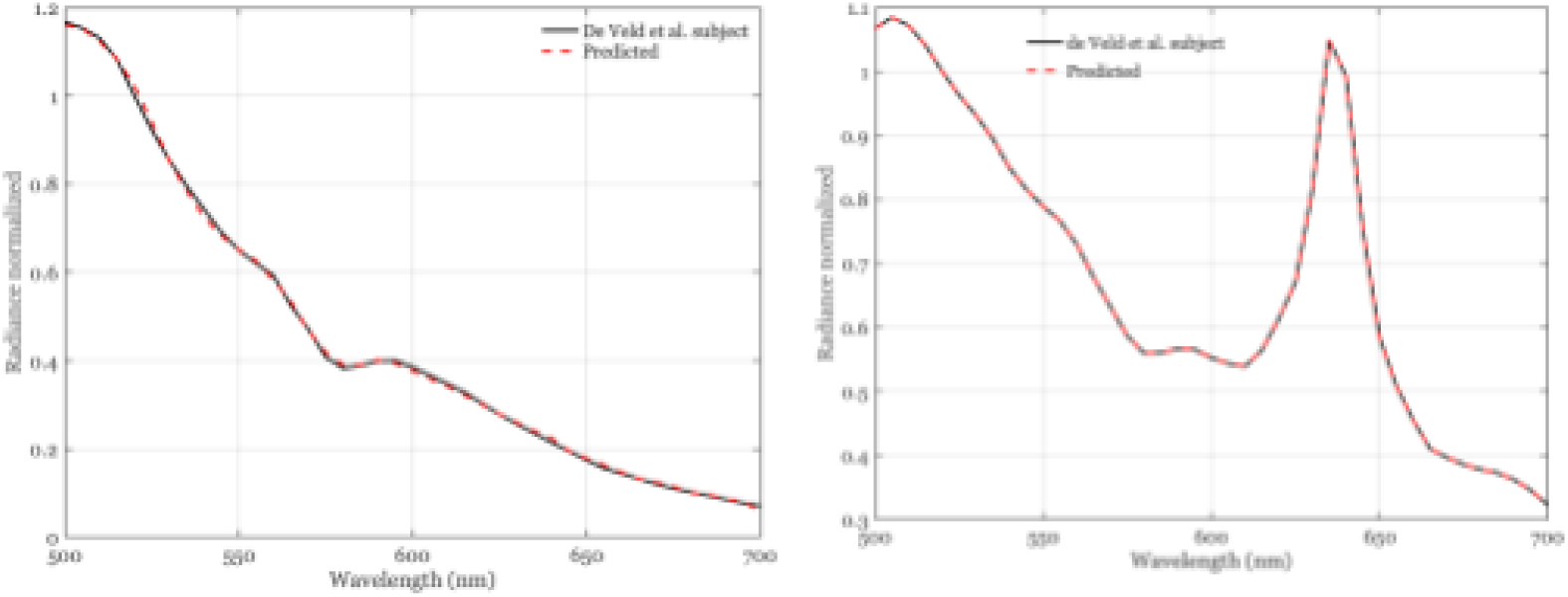
The lip (left) and tongue (right) measurements from a typical subject in (De Veld et al. 2003) are well approximated as the linear combination from the spectral fluorescence data in the present study. This agreement suggests that the same fluorophores are the basis of the independently measured data sets.

We conducted a similar analysis using data from Mallia et al. (Mallia et al. 2008) - who measured autofluorescence on the dorsal surface of the tongue with 404 nm excitation in 15 healthy volunteers - and from Betz et al. (Betz et al. 1999) - who illuminated the inner lip and dorsal surface of the tongue in 17 individuals with an excitation light that included wavelengths ranging between 375 and 440 nm. Our data could predict the mean autofluorescence spectra reported by these investigators for the dorsal tongue surface ((Mallia et al. 2008), (Betz et al. 1999)) and the inner lip ((Betz et al. 1999)) within 1-3%.

We are continuing to measure bulk fluorescence in healthy individuals to quantify the natural variability in blood density and fluorophore concentrations within the population. Expanding our database of spectroradiometric measurements will help establish reference values and population variance for bulk fluorescence in healthy tissue, which is important for developing systems that can differentiate between healthy and abnormal tissue.

## Conclusion

This study provides quantitative evidence supporting the use of autofluorescence imaging as an objective tool for assessing the health of oral mucosal tissue. By employing precise spectral radiance measurements and developing a simple tissue fluorescence model, we provide a method for estimating the relative contributions from several fluorophores, while also highlighting the critical role of components like blood attenuation in shaping these signals.

Building on prior work in digital imaging (Farrell et al. 2003; Farrell, Catrysse, and Wandell 2012; Farrell and Wandell 2015; Lyu et al. 2022), our team is developing a computational model referred to as a digital twin (Benson, 2023; Holopainen et al., 2022). Our digital twin integrates four critical elements: (1) 3D oral cavity geometry, (2) tissue spectral reflectance, (3) excitation-emission matrices for endogenous/exogenous fluorophores, and (4) camera optics/sensor characteristics (Farrell, J., Lyu, Z., Liu, Z., Blasinski, H., Xu, Z., Rong, J., Xiao, F., Wandell, B. 2019; Lyu et al. 2021). This digital twin enables systematic optimization of imaging components to isolate biological features like blood and fluorophores—a capability that could enhance detection of oral mucosal abnormalities through improved fluorescent signal differentiation.

To expand this framework, we will merge our quantitative autofluorescence model with AFI-system digital twins. This combined approach will permit virtual testing and optimization of excitation sources, optical filters, and sensor quantum efficiency under diverse biological conditions. By streamlining the evaluation of different configurations and predicting system performance prior to manufacturing, we aim to reduce reliance on physical prototypes and accelerate the development of advanced imaging systems for oral health monitoring.

## Acknowledgements

We gratefully acknowledge Henryk Blasinski, Haomiao Jiang, and Zhenyi Liu for their valuable feedback and suggestions on this manuscript. We also thank Doug Ward for his assistance with data collection and Krithin Kripakaran for his help in creating the illumination system.

## Footnotes

1 https://github.com/ISET/oe_tongue_lip

2 https://github.com/ISET/ISETCam/wiki

3 https://github.com/ISET/isetfluorescence/wiki

4 We modeled blood transmittance from the absorbance data in https://omlc.org/spectra/hemoglobin/summary.html.

5 The computations are in the script oeFigure5_9_ModelFits.mlx in https://github.com/ISET/oe_tongue_lip.

6 The original data are not available.

